# Load-Dependent Relationships between Frontal fNIRS Activity and Performance: A Data-Driven PLS Approach

**DOI:** 10.1101/2020.08.21.261438

**Authors:** Kimberly L. Meidenbauer, Kyoung Whan Choe, Carlos Cardenas-Iniguez, Theodore J. Huppert, Marc G. Berman

## Abstract

Neuroimaging research frequently demonstrates load-dependent activation in the prefrontal cortex during working memory tasks such as the N-back. Most of this work has been conducted in fMRI, but functional near-infrared spectroscopy (fNIRS) is gaining traction as a less invasive and more flexible alternative to measuring cortical hemodynamics. Few fNIRS studies, however, have examined how working memory load-dependent changes in brain hemodynamics relate to performance. The current study employs a newly developed and robust statistical analysis of task-based fNIRS data in a large sample, and demonstrates the utility of data-driven, multivariate analyses to link brain activation and behavior in this modality. Seventy participants completed a standard N-back task with three N-back levels (N = 1, 2, 3) while fNIRS data were collected from frontal and parietal cortex. Overall, participants showed reliably greater fronto-parietal activation for the 2-back versus the 1-back task, suggesting fronto-parietal fNIRS measurements are sensitive to differences in cognitive load. The results for 3-back were much less consistent, potentially due to poor behavioral performance in the 3-back task. To address this, a multivariate analysis (behavioral partial least squares, PLS) was conducted to examine the interaction between fNIRS activation and performance at each N-back level. Results of the PLS analysis demonstrated differences in the relationship between accuracy and change in the deoxyhemoglobin fNIRS signal as a function of N-back level in four mid-frontal channels. Specifically, greater reductions in deoxyhemoglobin (i.e., more activation) were positively related to performance on the 3-back task, unrelated to accuracy in the 2-back task, and negatively associated with accuracy in the 1-back task. This pattern of results suggests that the metabolic demands correlated with neural activity required for high levels of accuracy vary as a consequence of task difficulty/cognitive load, whereby more automaticity during the 1-back task (less mid-frontal activity) predicted superior performance on this relatively easy task, and successful engagement of this mid-frontal region was required for high accuracy on a more difficult and cognitively demanding 3-back task. In summary, we show that fNIRS activity can track working memory load and can uncover significant associations between brain activity and performance, thus opening the door for this modality to be used in more wide-spread applications.

## 1. Introduction

Functional near-infrared spectroscopy (fNIRS) is a neuroimaging modality that has gained traction in recent years due to its versatility to studying brain activity in realistic natural environments. Compared to electroencephalography (EEG) and functional magnetic resonance imaging (fMRI), fNIRS is robust to motion artifacts and environmental noise, making it an increasingly popular method for studying neural activity outside of standard laboratory experimentation (Pinti, Tachtsidis, et al., 2018; Yücel et al., 2017). fNIRS uses light spectroscopy at near-infrared wavelengths to measure the same cerebral metabolic changes that are measured using functional MRI (Buxton, 2010; Huppert et al., 2006). In both methods, the measurements taken are metabolic proxies for neuronal activity. When neural activity increases, so does the metabolic demand, leading to increased blood flow in the surrounding vasculature. This blood flow causes an increase in concentrations of oxygenated hemoglobin and a decrease in concentrations of deoxygenated hemoglobin (Buxton, 2013; Huppert et al., 2006).

Historically, many of the typical fNIRS study paradigms and analysis techniques mirrored those of task-based fMRI. The vast majority of existing fNIRS studies involved initial data preprocessing (i.e., downsampling, bandpass or wavelet filtering, motion correction), before being converted into oxyhemoglobin (HbO) and deoxyhemoglobin (HbR) concentrations (Huppert et al., 2009; Scholkmann et al., 2014). A general linear model was then applied to create contrasts in HbX changes between task conditions or between task and rest (Cooper et al., 2012; Pinti, Scholkmann, et al., 2018; Scholkmann et al., 2014). However, it was recently demonstrated that these typical approaches overall fail to account for specific statistical properties of the fNIRS signal, and in doing so, inflate the false positive rate of reported results (Barker et al., 2013; Huppert, 2016).

Due to these issues, new analysis methods have been developed to account for these specific statistical properties of fNIRS. By applying pre-whitening to the linear model to reduce noise correlations and using robust regression to down-weight statistical outliers, these methods perform better on sensitivity-specificity analyses and show better control of type-I errors (Barker et al., 2013; Huppert, 2016; Santosa et al., 2018). With proper statistical analysis to account for these unique noise properties, fNIRS provides an increasingly rigorous method of neuroimaging. Furthermore, the cost-effectiveness of fNIRS allows for larger sample sizes than in fMRI, lessening the risk of failures in replicability due to small samples (Turner et al., 2018).

One of the more well-studied effects in fMRI and fNIRS is that of cognitive load-dependent changes in frontal and parietal cortical regions (Cui et al., 2011; Fishburn et al., 2014; Herff et al., 2014; Mencarelli et al., 2019; Owen et al., 2005). That is, neural activity in these regions increases with more cognitively taxing and difficult tasks. Typically, this is achieved by increasing the number of items needed to be stored in working memory in a task requiring sustained attention, as in the N-back task (Conway et al., 2005; Kirchner, 1958; Owen et al., 2005). In an N-back task, participants are required to compare a current stimulus to a stimulus presented N items back during continuous presentation. This N may be manipulated (typical values include N = 0, 1, 2, or 3), thus indicating the number of items in working memory. While some previous fNIRS studies have shown linear increases in frontal activation based on N-back level (Ayaz et al., 2012; Fishburn et al., 2014; Kuruvilla et al., 2013), several others have found non-linear effects (i.e., activation that does not follow the pattern of 3-back > 2-back > 1-back > 0-back; Aghajani et al., 2017; Mandrick et al., 2016). In cases of non-linear increases in activation with greater task demands, researchers have proposed that participants may simply disengage from tasks that are too difficult (Causse et al., 2017). It has also been posited that once participants reach a maximum level of cortical activation during a demanding (but achievable) task, no additional neural “output” can be tapped into to perform well on an even more difficult task (Mandrick et al., 2013).

While these discrepancies are attributed to performance-based limitations on very demanding tasks, not all studies have attempted to explicitly link the load-dependent activation results to task accuracy, and those which did have yielded mixed results. When examined across a broader array of cognitive tasks which manipulate difficulty, some studies have not found significant correlations between fNIRS activity and performance in the cortical regions of interest (Ayaz et al., 2012; Causse et al., 2017; Matsuda & Hiraki, 2006). Other work has identified negative relationships between task performance and cortical activation when participants undergo working memory training, resulting in increased neural efficiency (McKendrick et al., 2014). Results of one fNIRS study examining the role of expertise on prefrontal activity and task difficulty suggested that this relationship is a complex one, but did not directly link activation to behavior (Bunce et al., 2011). Ultimately, while there is evidence that performance influences task-evoked fNIRS activity, how exactly this relationship is affected by task demands remains an open question.

The primary goal of the current study was to conduct a well-powered validation of the load-dependent fNIRS responses demonstrated in prior work using a traditional verbal N-back task, large sample, and utilizing recently developed robust statistical analytical procedures. It was hypothesized that during the N-back task, prefrontal and parietal cortical activity would be largest for the 3-back task (highest cognitive load), lessened for the 2-back task (medium cognitive load), and smallest for the 1-back task (lowest cognitive load).

A second, exploratory goal was to examine how individual differences in participant accuracy could affect load-dependent fNIRS activity. This work sought to examine whether the relationship between fNIRS activity and performance differed based on task difficulty, specifically by using a data-driven, multivariate partial least squares (PLS) analysis to evaluate this relationship. Behavioral PLS (McIntosh & Lobaugh, 2004) has been used in other neuroimaging modalities (such as fMRI, EEG, and magnetoencephalography (MEG)) as a data-driven approach to extract relationships between neural activity and behaviors of interest (Bialystok et al., 2005; Chang et al., 2017; Krishnan et al., 2011; Lobaugh et al., 2001; McIntosh et al., 2008), but has not yet been implemented in fNIRS research. Therefore, the current study also tested the utility of a multivariate PLS approach in fNIRS research to shed light on how the link between performance and neural activation may be affected by task demands.

In summary, the current study was designed to replicate the effect of load-dependent activation in frontal and parietal cortex in fNIRS using more robust statistical analyses. As with several previous studies, we found non-linear activation effects, which are likely attributable to poor performance on the 3-back task. To better elucidate the impact of performance on activation as a function by N-back level, a behavioral PLS analysis was conducted, and provided insight into how the relationship between accuracy and prefrontal activation differs based on task difficulty.

## 2. Method

### 2.1 Participants

Seventy adults participated in this study. All participants had normal or corrected-to-normal visual acuity. Participants gave written informed consent before participation and experimental procedures were approved by the University of Chicago’s Institutional Review Board (IRB). Participants were compensated $26 or 2 units of course credit, plus a performance-based bonus of up to $10. The full procedure included additional study elements related to a video intervention that were separate from the current work and lasted approximately 15 minutes. The total duration of the study was between 75 and 90 minutes.

Two participants were excluded from all data analysis due to participant non-compliance with the study procedures. Six additional participants were excluded from fNIRS analysis due to technical issues (2 participants) or low quality fNIRS data (4 participants), leaving a final sample of 62 participants. Of the 62 participants with usable fNIRS data, 28 were male and 34 were female, and the mean age was 23.6 years (*SD* = 6.3 years).

### 2.2 fNIRS Data Acquisition

fNIRS data were collected from a continuous-wave NIRSport2 device (NIRx Medical Technologies, LLC). The wavelengths of emitted light (LED sources) in this system were 760 nm and 850 nm, corresponding to oxygenated hemoglobin and deoxygenated hemoglobin concentrations, respectively. The data were collected at a sampling rate of 4.5 Hz using the NIRx acquisition software, Aurora fNIRS. The fNIRS cap contained a total of 16 sources and 16 detectors creating 43 total channels covering bilateral frontal cortex (33 channels) and right parietal cortex (10 channels).

### 2.3 fNIRS Optode Locations (Montage)

The montage was created using fOLD (fNIRS Optodes’ Location Decider; Morais et al., 2018), which allows placement of optodes in the international 10-10 system to maximally cover anatomical regions of interest, as specified by one of 5 parcellation atlases. The AAL2 (Automated Anatomical Labeling; Rolls et al., 2015) parcellation was used to generate the montage, which was designed to provide as much coverage of the prefrontal cortex (PFC) as possible, covering bilateral superior and inferior frontal gyri. This emphasis on frontal cortical areas was decided based on evidence from other N-back studies using fMRI (see Owen et al., 2005 for a meta-analysis) and fNIRS, which have demonstrated that load-dependent changes in HbO and HbR are found across areas of the PFC (Ayaz et al., 2012; Fishburn et al., 2014; Herff et al., 2014; Sato et al., 2013).

The right parietal region was selected as an additional ROI for this task due to evidence that parietal cortical regions are engaged during attention-demanding tasks in fNIRS (Hosseini et al., 2017; Murata et al., 2015). As parietal data quality is usually less consistent than channels unobstructed by hair (such as the forehead), the majority of optodes (12 sources and 12 detectors) were focused on prefrontal regions, leaving only 4 sources and 4 detectors to cover parietal areas. Rather than sparsely covering bilateral parietal cortex, better coverage of right parietal cortex was examined in the current study. Right parietal was chosen as participants would be required to use their right hand to respond (hence activating left motor/sensorimotor areas) during the task and our parietal montage overlapped with the standard sensorimotor fNIRS montage. As we did not want to have the more anterior channels in our parietal montage to be affected by differences in contralateral sensory or motor-evoked activity (i.e. due to less or more responding based on task difficulty), we opted to focus on right parietal coverage. While verbal working memory storage and rehearsal are more associated with left-lateralized regions of parietal cortex (Awh et al., 1996; Ravizza et al., 2004), some meta-analytic data demonstrate bilateral parietal activation across verbal and non-verbal N-back tasks (Mencarelli et al., 2019; Owen et al., 2005). [**Fig. 2**]

Gross ROIs from the montage (used in subsequent figures) were defined based on the Brain AnalyzIR Toolbox’s depth map function (Santosa et al., 2018). Depth maps show the distance from each fNIRS optode to the superficial cortex of several talairach daemon labeled regions of the Colin27 atlas (Lancaster et al., 2000), which can be used to determine coverage of an ROI based on the montage used. As a topological fNIRS layout cannot access depths greater than approximately 30 mm, the channels (lines) projected over yellow or orange regions in **Fig. 3** (representing depths > 30 mm) are ones that do not reach the specified ROI, whereas channels covering green or blue areas are within range of the nearest cortical point within the ROI.

### 2.4 Procedure

After providing informed consent, experimenters measured the participants’ head to determine cap size and placement, then began to set up the cap while participants were taken through task instructions and given an opportunity to practice the N-back task. After the first round of practice, the cap was placed on the participants' head, moving hair as needed to provide clear access to the scalp for the sources and detectors. Cap alignment was verified based on the international 10-20 location of Cz (Klem et al., 1999). fNIRS data were then calibrated and checked for quality before proceeding. If any channels were not displaying sufficiently high quality data, placement and hair-clearing were performed again before continuing. Next, participants completed a short round of additional practice (single block of each 1-back, 2-back, and 3-back without trial-by-trial feedback), before continuing to the main round of the N-back task. After completing the experiment, the cap was removed and participants completed a demographics questionnaire. All experimental procedures were coded and presented using PsychoPy (Peirce et al., 2019).

### 2.5 N-back Task

The experimenter took participants through step-by-step instructions of the N-back task before participants began practice. Participants were told that they would see a sequence of short words that are separated by brief fixations, and that every 2 seconds a word would be presented that should be compared to the word “N” trials back. In the current study, N was 1, 2, or 3. Participants were instructed to press the “m” key every time the current word matched the word N trials back, and to press the “n” key every time the current word did not match the word N trials back **[Fig. 1]**. Each block began by displaying the N-back level and a fixation cross (5 seconds). Each task block contained a 15-length pseudorandom sequence of two words, presented for 2 seconds each for a total of 30 seconds, followed by 20 seconds of rest. Therefore, the length of each block was 55 seconds. To suppress sequence memory formation, the two words used in each block were randomly selected from the eight word pool (‘WHAT', 'HOW', 'WHEN', 'WHY', 'WHERE', 'WHO', 'THAT', 'BUT'), except during the first practice, in which the words “AXE” and “BOX” were used. In addition, the sequence of two words was determined using an m-sequence (base = 2, power = 4; thus one word appeared eight times, and the other word appeared seven times; Buračas & Boynton, 2002; Choe et al., 2014; Choe et al., 2016) to suppress its autocorrelation. In all cases, words were presented in white text on a black background.

**Fig 1.**
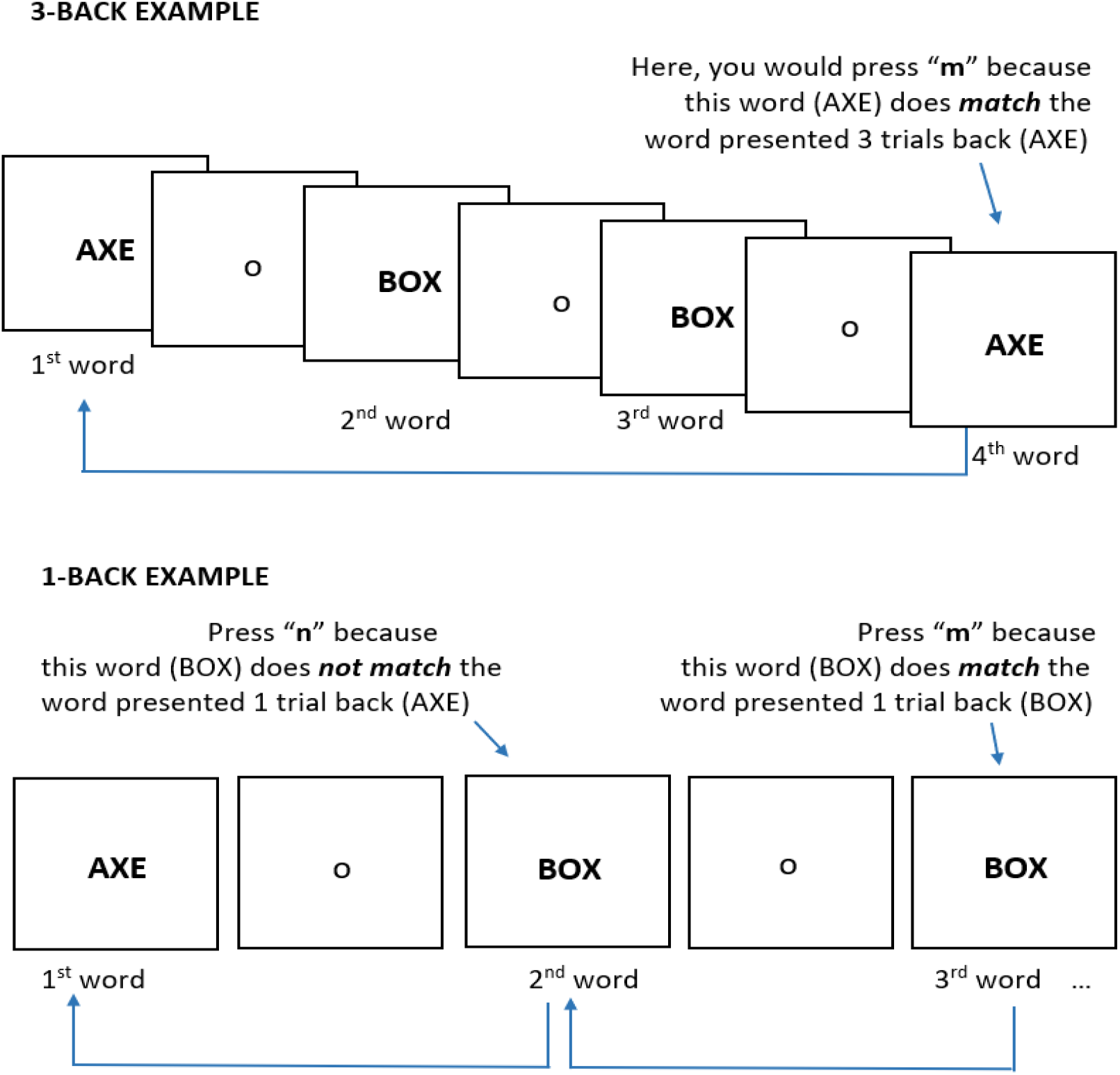
N-back Task. Example of 1-back task (Top) and 3-back (Bottom). 2-back task not shown.

**Figure 2.**
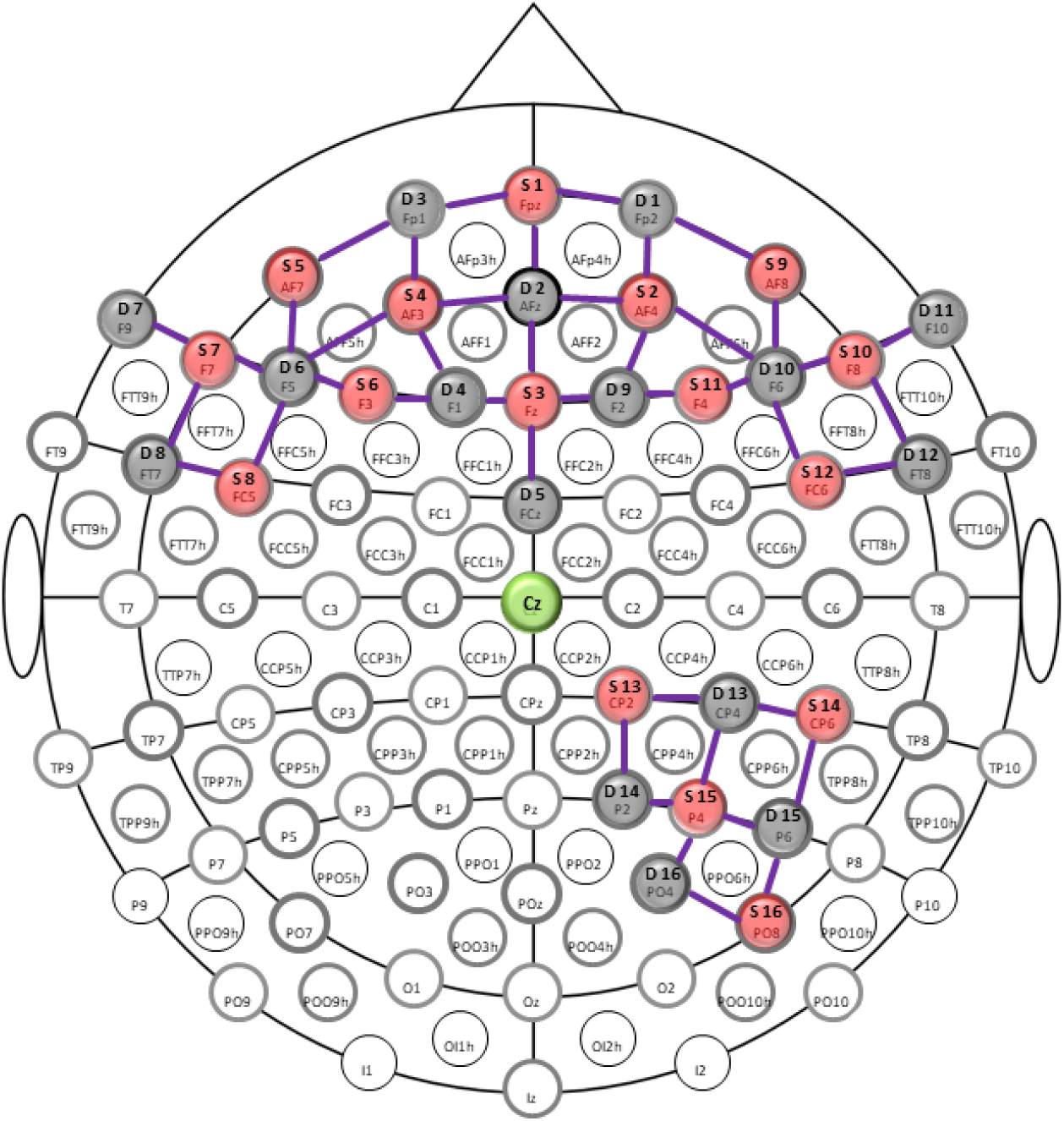
fNIRS Montage in international 10-10 coordinate space. Montage with 16 x 16 frontal source-detector pairs and 4 x 4 right parietal source-detector pairs. Sources are indicated in red, detectors are indicated in gray, and channels are indicated by purple lines. Cz highlighted in green.

**Figure 3.**
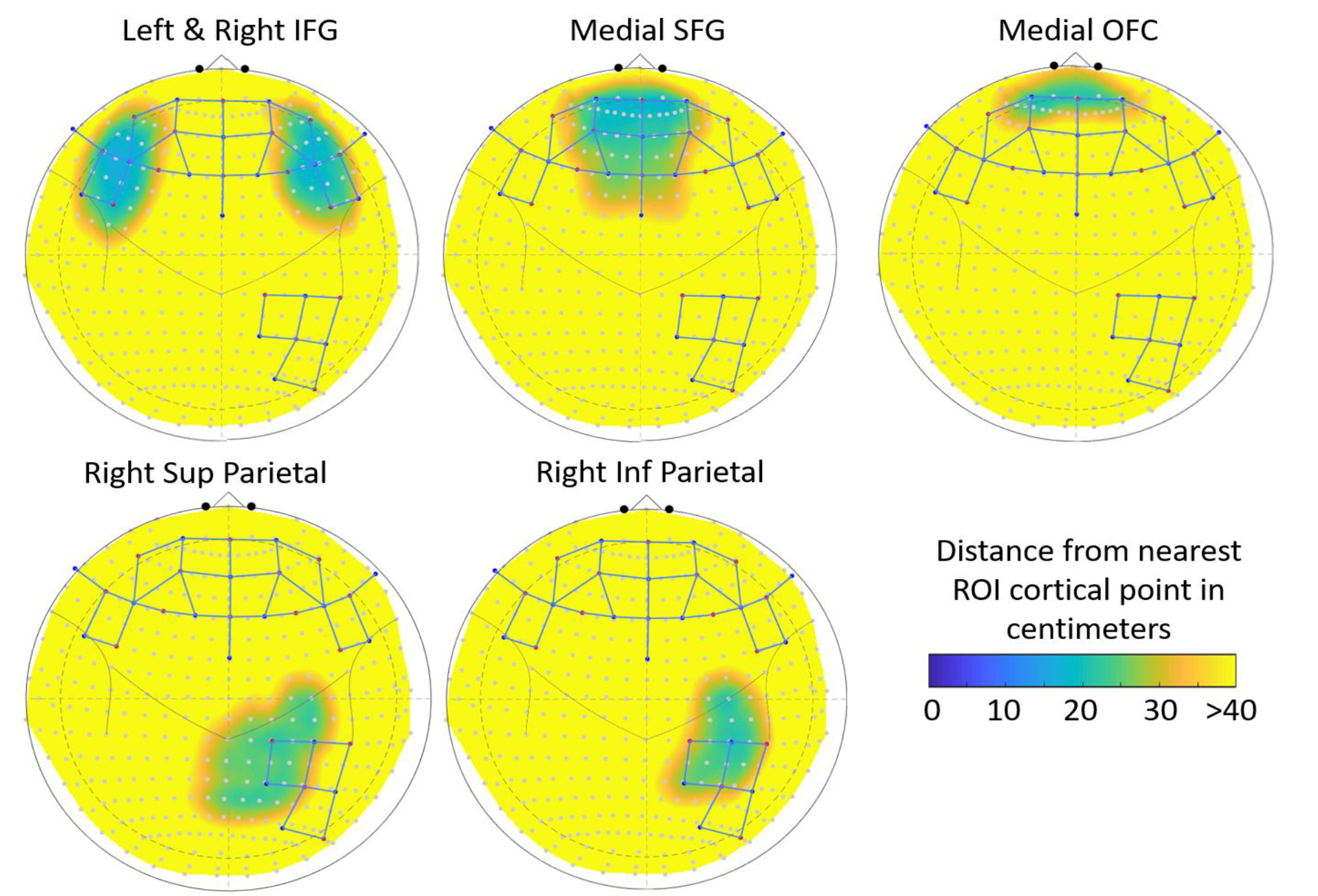
Gross ROI depth maps with superimposed montage. fNIRS montage (registered to Colin27 atlas) and depth map for 6 ROIs taken from the talairach daemon parcellation: Left and Right Inferior Frontal Gyrus, Medial Superior Frontal Gyrus, Medial Orbitofrontal Cortex, Right Superior Parietal Gyrus, Right Inferior Parietal Gyrus.

After the experimenter took participants through the N-back instructions participants performed the first round of N-back practice, consisting of 9 blocks. In this first practice, accuracy feedback was provided on a trial-by-trial level as well as at the end of each block. Participants completed 3 blocks of 1-back, then 3 blocks of 2-back, and then 3 blocks of 3-back. After the fNIRS cap was set up, participants began the second round of practice designed to more closely mimic the conditions of the real task. In this practice, participants performed a single block of 1-back, then 2-back, then 3-back, without trial-by-trial feedback. The main N-back task involved 18 blocks, with 6 blocks of each N-back level, pseudorandomly presented.

Participants received a performance-based bonus during this round of N-back task, wherein performance > 90% on a block earned an additional 40 cents per block, > 80% earned an additional 30 cents per block, and > 60% earned an additional 20 cents per block. Performance under 60% did not yield a cash bonus in this study.

### 2.6 Analysis

#### 2.6.1. Behavioral Analysis

Accuracy on the N-back task was calculated by taking the average accuracy over the 6 blocks of each N-back level. Statistical analysis was conducted using R version 3.5.1 (R Core Team, 2018). Accuracy-level differences between levels of the main N-back task, were analyzed with a repeated measures ANOVA using function ‘ezANOVA’ in the ‘ez’ package (Lawrence, 2016), effect size was calculated using the function ‘eta_sq’ in the ‘sjstats’ package (Lüdecke, 2020), and Bonferroni corrected post-hoc contrasts were conducted using paired t-tests in the R ‘stats’ package.

#### 2.6.2. fNIRS Data Analysis: Quality Check

fNIRS data were first loaded into the HOMER2 software package (Theodore J. Huppert et al., 2009) for visual inspection and segmentation of the main N-back trials from practice trials. Visual inspection was done to examine overall data quality (at the level of the participant) and to assess the quality of the parietal data, which was much more noisy and variable than the frontal data. Visual inspection was performed by examining the power spectral density plots for all channels to identify the presence of a cardiac oscillation, which is typically around 1 Hz (Tong et al., 2011). The presence of this cardiac signal is a good indicator that the optical density signals are successfully coupled with a physiological hemodynamic response (Hocke et al., 2018). This method was used to do a first pass evaluation. Based on this visual inspection, 4 participants with unusable data (defined as 5 or fewer clean channels) were identified and excluded from further analyses. Parietal data quality was also examined and logged to determine whether analysis of this region would be fruitful. Of the 62 participants that were kept, 17 had fully usable parietal data, 20 had mostly usable parietal data (at least half of channels showing good physiological coupling), and 25 had unusable parietal data (only a few usable channels or none). It should be noted that, as the statistical analysis downweights noisy channels in the linear model (see next section), including these channels will not increase the likelihood of a false positive effect, but the power to detect an effect in this area is reduced.

#### 2.6.3. fNIRS Data Analysis: Pre-processing Pipeline and Task-Based Activation

fNIRS data were then analyzed using the NIRS Brain AnalyzIR Toolbox (Santosa et al., 2018). Using this toolbox, the.nirs data (raw light intensity) were loaded into the program, converted into optical density, then converted to oxygenated (HbO) and deoxygenated (HbR) hemoglobin concentrations using the modified Beer-Lambert law (Strangman et al., 2003).

Once the data were in the form of HbO and HbR concentrations, first level (subject-level) statistics were calculated. As alluded to previously, fNIRS data have unique statistical properties that are not accounted for by typical fMRI-based analysis, and can inflate the type-I error rate (Huppert, 2016). In particular, unlike fMRI, fNIRS suffers from serially-correlated errors (due to a higher sampling rate than the physiological signal of interest) and heavy-tailed noise distributions (due to motion-related artifacts and often, large differences in SNR between channels and between participants; Huppert, 2016). To correct for these issues, the first level general linear model run on individual participants’ data uses an autoregressive, iteratively reweighted least-squares model (AR-IRLS). The AR-IRLS model employs an auto-regressive filter (pre-whitening) to deal with the serially correlated errors and uses robust weighted regression to iteratively down-weight outliers due to motion artifacts (Barker et al., 2013). This model saves both the subject level regression coefficients and their error-covariance matrices to be used in statistical tests and contrasts for each subject, and eventually, for use in second-level (group-level) analyses.

Based on research investigating the sensitivity and specificity of basis sets in fNIRS as a function of signal quality and task period (Santosa et al., 2019), a canonical HRF basis set was selected for this analysis. Work by Santosa et al. (2019) found that for tasks of sufficiently long durations (>10 seconds, as in the current study), the canonical HRF performs best in a sensitivity-specificity (ROC) analysis. The canonical model has lower degrees of freedom than a full deconvolution of the raw hemodynamic response (finite impulse response, or FIR model), which improves performance on ROC analysis. This is true at durations of more than 10 seconds, even though there may be a mismatch between the shape of the canonical HRF and the actual hemodynamic response (Santosa et al., 2019).

Based on the output of the first level statistical models, 3 subjects with undue leverage for the group analyses were calculated (those which contribute significant leverage towards the group results, defined by subject-level leverage of *p* < 0.05) and were removed from group-level analyses. Next, second-level statistical models were calculated, which use the full covariance from the first-level models to perform a weighted least-squares regression (Santosa et al., 2018). Robust regression was also applied to the second-level model to down-weight outliers at the group-level. The results of this analysis were used for group-level contrasts between N-back levels at each channel.

Group activation results are reported as statistical maps using Benjamini-Hochberg false-discovery rate-corrected p-values (e.g., *q*-values; Benjamini & Hochberg, 1995). This FDR correction was applied to all of the data in the second-level analysis, including 43 channels, oxy- and deoxy-hemoglobin, and 3 conditions, making the correction very conservative over all tests.

#### 2.6.4. fNIRS Data: Behavioral PLS Analyses

Behavioral PLS analysis (Berman et al., 2014; McIntosh & Lobaugh, 2004; https://www.rotman-baycrest.on.ca) was conducted to identify significant relationships between fNIRS activity and task performance as a function of N-back level. PLS (partial least squares) analysis is a multivariate, data-driven approach often used to examine brain-behavior associations in neuroimaging research by relating two sets or “blocks'' of data to one another (Krishnan et al., 2011). In this study, the fNIRS data block consisted of the regression coefficients (*ꞵ*) from the first-level statistical model (AR-IRLS), corresponding to changes in HbO or HbR for each N-back level relative to baseline for each participant. The behavioral block consisted of average accuracy for each N-back level across blocks of the main N-back task for each participant. Thus, for each PLS (HbO or HbR), each participant had 129 values for the brain activity block (activation betas for each of three N-back levels for 43 channels) and three values for the behavioral block (average accuracy for each of three N-back levels). The goal of this analysis was to find the linear combination of conditions and brain activity that maximized their covariance. These weighted patterns are referred to as latent variables (LVs). Behavioral PLS is a variant of PLS that examines brain-behavior relationships as a function of condition. The resulting behavioral PLS analysis identified the correlation between behavioral data (in this case accuracy) and brain activity by condition type that again maximized the covariance between brain activity and conditions. Therefore, in this behavioral PLS, the LVs represent a specific pattern of brain-performance relationships that vary by condition.

Before running the PLS analysis, histograms of the fNIRS beta values were plotted to examine whether the brain data contained any extreme outliers that may bias the PLS analysis and would ordinarily be removed in the AnalyzIR Toolbox’s robust regression (Huppert, 2016). One participant contained extreme outliers at channel 29 (i.e., beta values < −100 and > 100), and was therefore excluded from PLS analysis. However, the direction and significance of results did not change if this participant was included. Ten thousand permutation tests were performed to obtain p-values for each latent variable (LV) and 10,000 bootstrapped samples with replacement were created to generate the 95% confidence intervals for the mean correlation between fNIRS activity and performance by condition for each channel. The bootstrap ratios (salience[weights]/SE[reliability]) measure the reliability of the brain-behavior relationship at each channel, and a larger bootstrap ratio indicates a strong and consistent contribution to the LV. In this study, channels with bootstrap ratios larger than +3 or smaller than −3 were determined to be statistically significant as these bootstrap ratios can be interpreted as z-scores.

### 2.7 Data & Code Availability

Data, analysis code, results, and experiment code are publicly available at: https://osf.io/sh2bf/

## 3. Results

### 3.1 Behavioral Results: N-back Performance

Results of the repeated measures ANOVA examining accuracy as a function of N-back level in the main task yielded a significant effect of N-back level on accuracy, *F*(2,120) = 80.0, *p* < 0.001, *ηp*^*2*^ = 0.57, 95% CI [0.45, 0.65]. As expected, accuracy for the 1-back task (*M* = 0.91, *SD* = 0.09) was significantly better than accuracy for the 2-back task (*M* = 0.79, *SD* = 0.17, *p* < 0.001) and for the 3-back task (*M* = 0.73, *SD* = 0.15, *p* < 0.001). Accuracy for the 2-back task was also significantly higher than for the 3-back task (*p* < 0.001). [Fig. 4]

**Figure 4.**
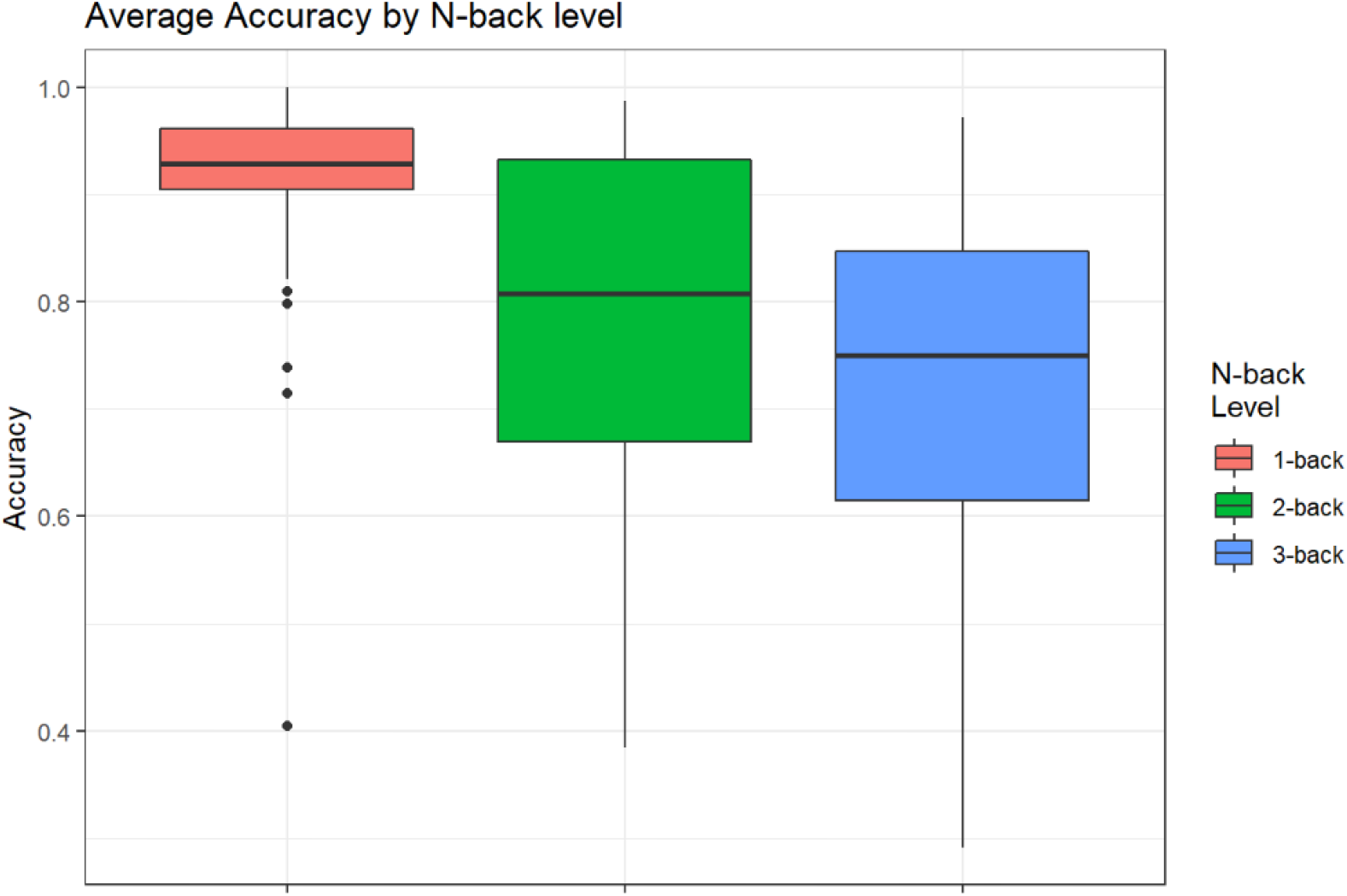
Boxplots of Average Accuracy by N-back Level for all participants

### 3.2 fNIRS Activation Results

#### 3.2.1 Activation vs. Baseline in Main N-back Task

In the GLM, baseline is defined by everything that is not a task event (i.e., is a test against the DC regressor in the model). Relative to baseline, significant increases in oxygenated hemoglobin (HbO) were found for 1 channel (medial superior frontal gyrus) for the 1-back task and for 5 frontal channels and 1 parietal channel for the 2-back task. No channels showed significant increases in HbO concentrations for the 3-back task. [**Table 1**] No channels showed significant decreases in HbR for any N-back level.

**Table 1.** Significant Activation by Channel & ROI for each N-back level. Significant channels (Source (S) - Detector (D) pairs) identified as FDR-corrected *q* < 0.05. ROI defined by maximal coverage of talairach daemon parcellation ROI. *p*-value listed is before FDR correction. Power listed is the estimated type-II power for that entry (calculated by computing the minimum detectable change as detailed in Harcum & Dressing, 2015).

|  | S | D | ROI | *t-stat* | *p* | *q* | *power* |
| --- | --- | --- | --- | --- | --- | --- | --- |
| **1-back HbO** | 1 | 2 | L Medial Superior Frontal Gyrus | 3.24 | 0.001 | 0.041 | 0.77 |
| **2-back HbO** | 2 | 1 | R Superior Frontal Gyrus | 3.48 | 0.001 | 0.032 | 0.83 |
|  | 4 | 2 | L Superior Frontal Gyrus | 3.28 | 0.001 | 0.042 | 0.78 |
|  | 4 | 3 | L Superior Frontal Gyrus | 3.66 | < 0.001 | 0.023 | 0.88 |
|  | 2 | 10 | R Middle Frontal Gyrus | 3.3 | 0.001 | 0.042 | 0.79 |
|  | 11 | 10 | R Inferior Frontal Gyrus | 3.86 | < 0.001 | 0.017 | 0.91 |
|  | 15 | 15 | R Inferior Parietal | 4.41 | < 0.001 | 0.004 | 0.97 |
| **3-back HbO** | - | - | - | - | - | - | - |

#### 3.2.2. fNIRS Contrasts: 2-back vs. 1-back

For HbO, 18 channels in the bilateral frontal and right parietal cortex showed significantly larger (*q* < 0.05) increases during the 2-back task relative to the 1-back task. No channels yielded larger HbO increases for the 1-back task relative to 2-back.

For HbR, 7 channels, primarily in bilateral inferior frontal gyrus (IFG), displayed larger decreases for 2-back over 1-back. Additionally, 5 channels, primarily in the right middle occipital gyrus, yielded larger decreases in HbR for 1-back relative to 2-back. [**Fig. 5, Top Panel**]

**Figure 5.**
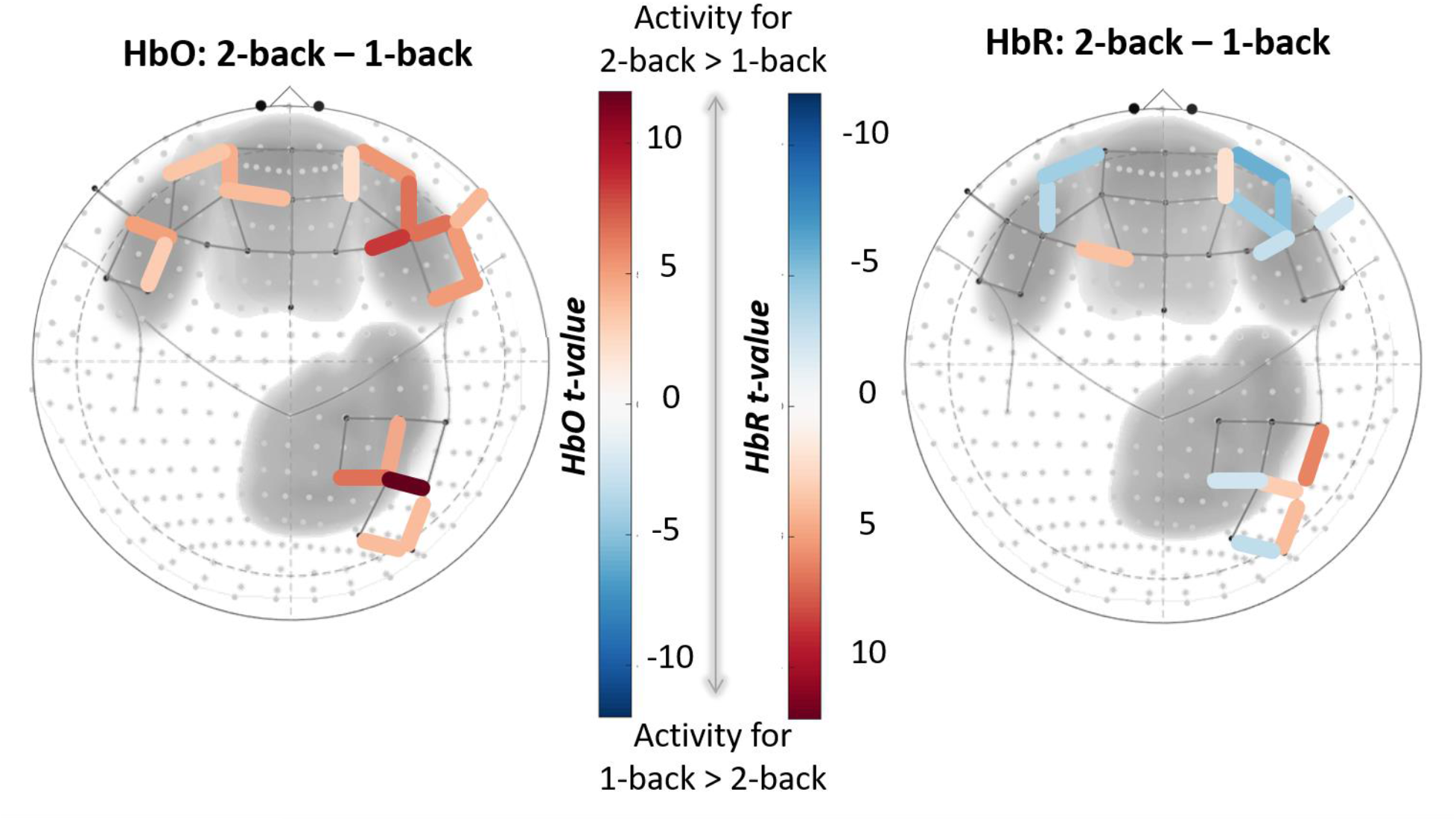
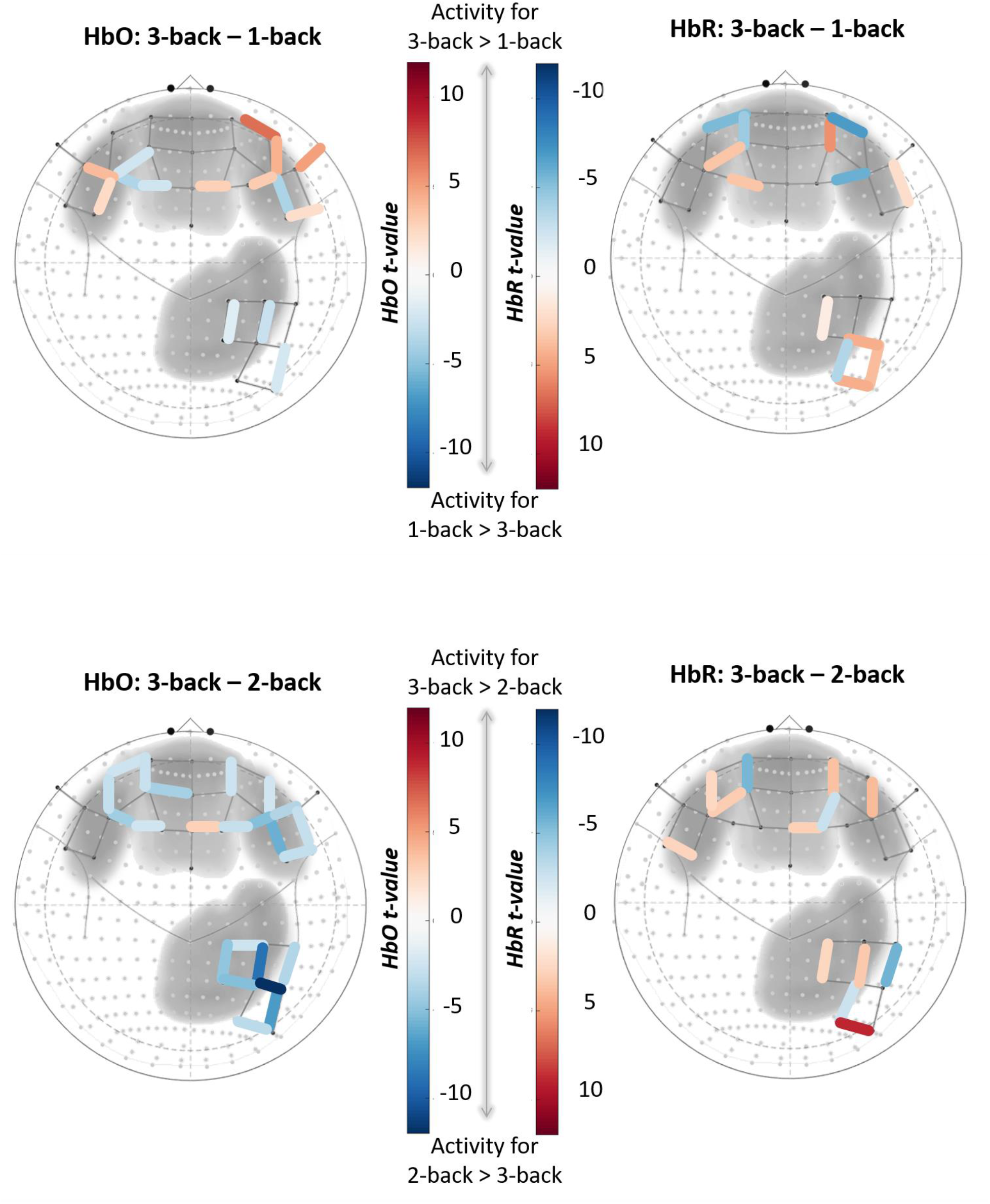
N-back level contrasts for HbO (left) and HbR (right). Only significant channels (*q* < 0.05) are shown. Channels are displayed on top of 10-20 coordinates and grayscale depth maps for left and right Inferior Frontal Gyri, medial Superior Frontal Gyri, and right Superior and Inferior Parietal Gyri. For HbO contrasts, positive *t*-values (red) correspond to relatively larger activity for the first term in the contrast, and negative *t*-values (blue) correspond to larger activity for the second term. The opposite pattern applies to HbR contrasts.

#### 3.2.3. fNIRS Contrasts: 3-back vs. 1-back

For HbO, 8 channels, primarily in left and right IFG, yielded significantly larger increases for 3-back relative to 1-back. Larger HbO increases for 1-back over 3-back were found in 7 channels, primarily located in the right inferior parietal cortex and left superior frontal gyrus (SFG).

For HbR, 5 channels (4 prefrontal, 1 inferior parietal), demonstrated larger deactivation in the 3-back task compared to 1-back. Eight channels (4 frontal and 4 occipito-parietal) showed the opposite pattern. [**Fig. 5, Middle Panel**]

#### 3.2.4. fNIRS Contrasts: 3-back vs. 2-back

For HbO, 22 channels showed significantly larger increases during the 2-back task compared to the 3-back task. These channels covered bilateral frontal and right parietal areas. Only one frontal channel was greater for the 3-back relative to the 2-back task.

For HbR, 9 channels distributed across bilateral frontal and right parietal cortex showed larger decreases for the 2-back task relative to the 3-back, and 4 channels (2 in medial SFG and 2 in inferior parietal cortex) showed the inverse pattern. [**Fig. 5, Bottom Panel**]

#### 3.2.5. fNIRS Contrasts Summary

Group level activation maps and contrasts between N-back conditions showed the most consistent results in the 2-back task relative to baseline and comparing activity during the 2-back task relative to the 1-back task. The consistently higher HbO and lower HbR concentration changes during the 2-back task, but not 3-back task, suggest that a minimum level of accuracy may be needed to elicit reliable activation in the fronto-parietal cortical regions examined. By minimum accuracy, we mean that if participants are not performing with a high enough accuracy they may not actually be engaged in the task because it has become too difficult. Participants overall performed more poorly on the main 3-back task. For the 59 participants used in group-level analysis of the main N-back task^1^, the average accuracy was 73.6% for the 3-back task. In comparison, average accuracy for these 59 participants was 80% for the 2-back task and 92.3% for the 1-back task.

### 3.3. Behavioral PLS Analysis: fNIRS Activity ~ Task Performance

Separate behavioral PLS analyses were run to relate performance to concentration changes in HbO and HbR by condition (i.e., 1-back, 2-back and 3-back). Though no statistically significant LVs were found for oxyhemoglobin (HbO), the first latent variable from the analysis with deoxyhemoglobin concentrations (HbR) was significant and explained 51% of the crossblock covariance (*p* = 0.025). Four superior frontal gyrus (SFG) channels (#4, #8, #12, and #25) showed N-back level dependent changes in the relationship between HbR and task accuracy [**Table 2**]. All of these significant channels had bootstrap ratios > 3, indicating the direction of the brain-behavior relationship was the same across all four channels. Specifically, for these channels, a larger reduction in HbR (equivalent to increased neural activity) was positively correlated with higher performance on the 3-back task, unrelated to activity on the 2-back task, and negatively correlated with performance on the 1-back task. In summary, this suggests that the metabolic demands placed on the prefrontal cortex that are necessary to achieve a high level of accuracy varies as a consequence of how difficult the task is. [Fig. 7]

**Table 2.**
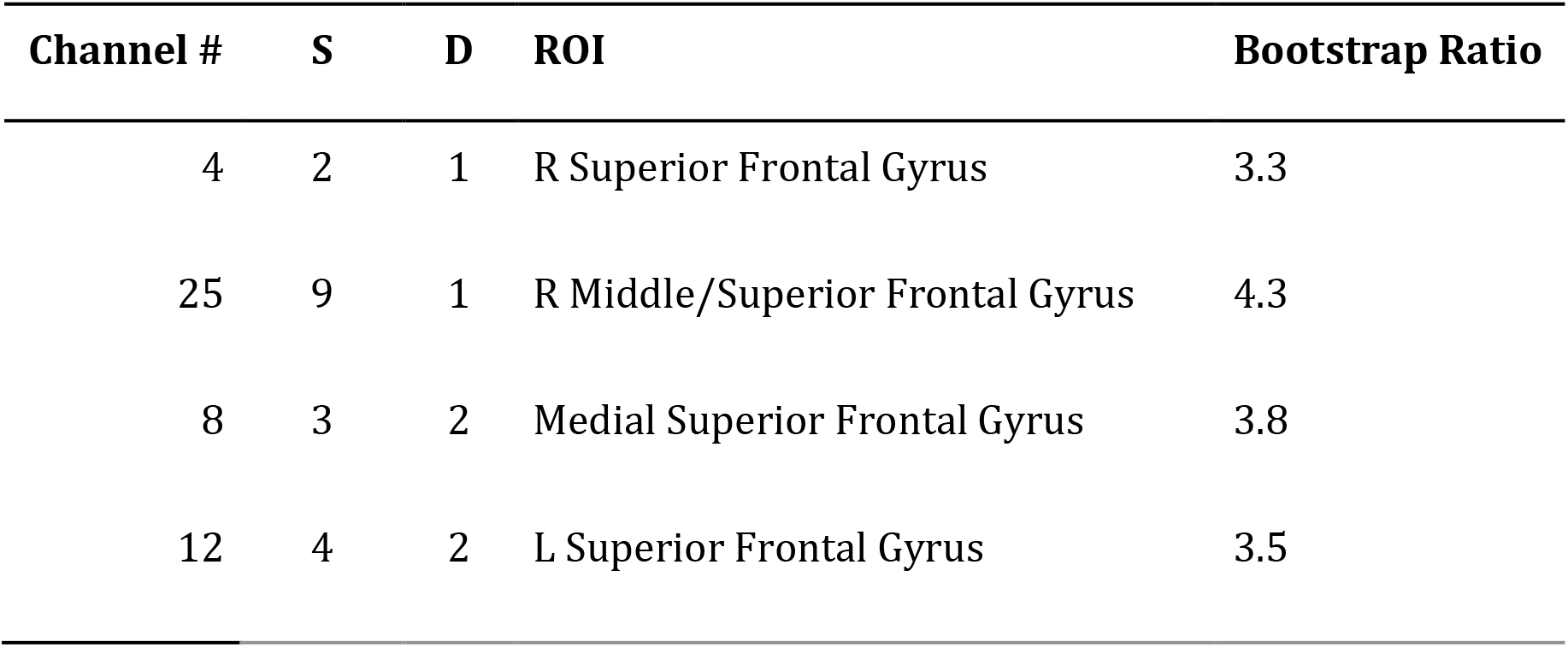
Significant Channels for LV 1 Channel number based on source (S) - detector (D) pair. ROI label defined by maximal coverage of talairach daemon ROI. Bootstrap ratios > 3 were considered significant.

**Figure 7.**
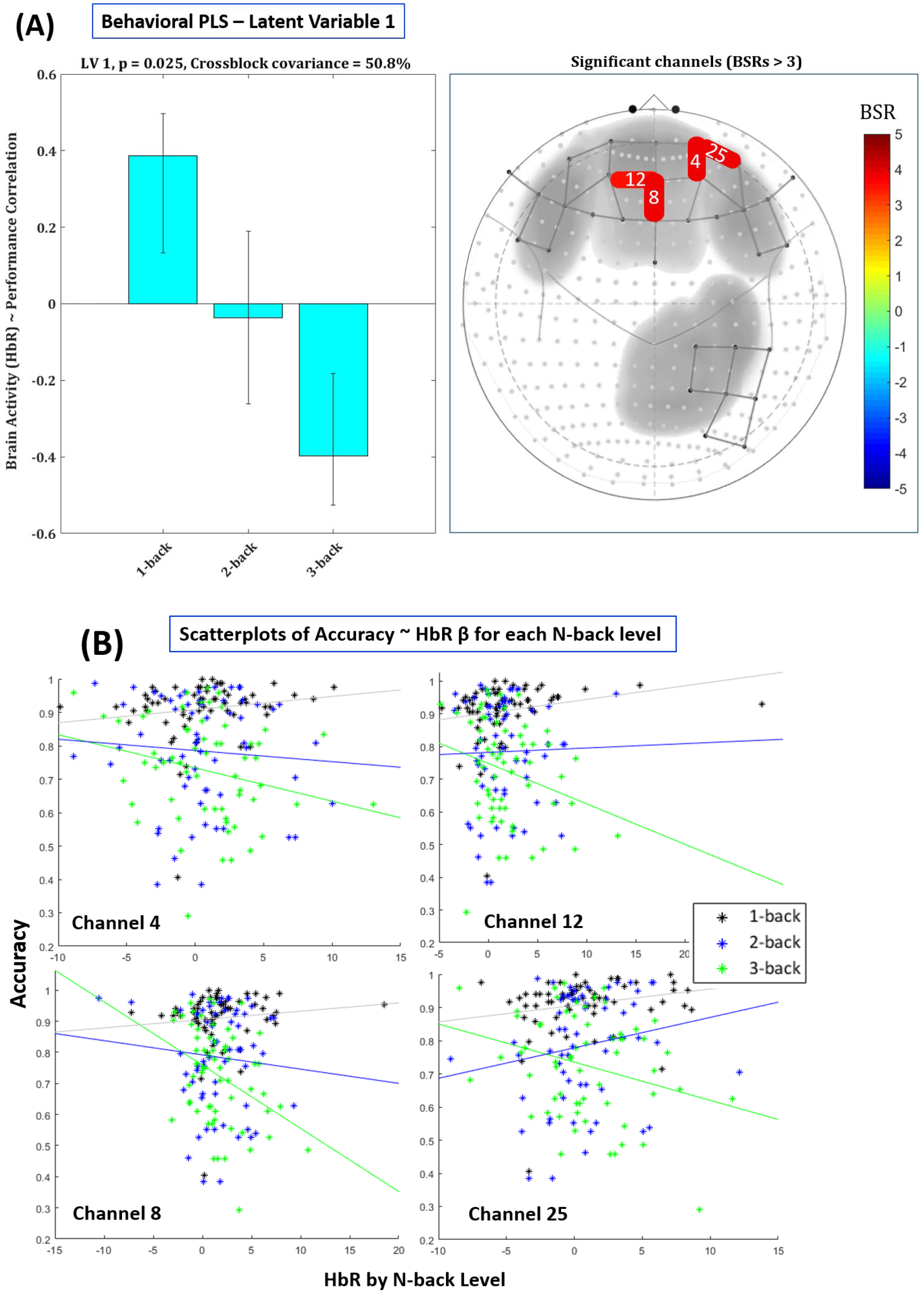
Results from first latent variable for HbR. LV 1 demonstrated an N-back load-dependent relationship between changes in deoxyhemoglobin concentrations (HbR) and performance. (**A**) The left panel shows correlation between accuracy and HbR concentration change separately by N-back level. Error bars are 95% confidence intervals around the mean correlation. The right panel shows significant channels (labeled by number), which had bootstrap ratios (BSR) > 3. (**B**) Scatterplots showing the correlation between HbR (*ꞵ* for task-evoked change from baseline) and performance (accuracy) at each channel, separated by N-back level.

## Discussion

The initial, confirmatory aim of this study was to further validate the use of fNIRS for measuring cognitive load with a large sample and utilizing recently developed robust statistical tools. Though a number of previous fNIRS studies have examined prefrontal activity using attention demanding working memory tasks such as the N-back (Aghajani et al., 2017; Ayaz et al., 2012; Fishburn et al., 2014; Kuruvilla et al., 2013; Sato et al., 2013), recent work has demonstrated that due to the unique statistical properties of fNIRS, the standard analysis approach (based on fMRI) can severely inflate the false positive rate (Huppert, 2016). In addition, discrepancies between studies that may be related to task performance have been demonstrated across a number of studies. Therefore, to provide convergent evidence for previous studies examining load-dependent changes in PFC and parietal cortex, a standard N-back task was employed with a large sample of participants, and using the recently developed Brain AnalyzIR Toolbox (Santosa et al., 2018) to deal with these fNIRS-specific statistical properties. As an exploratory aim, a behavioral PLS analysis was conducted to directly examine how performance and activation might be dependent upon cognitive load, which may explain non-linear load effects in this study and in previous work.

Overall, the fNIRS results were consistent with the general hypothesis that tasks placing higher demands on attention and working memory would lead to increased frontal and parietal activation as measured by HbO and HbR concentration changes. This was most evident by the widespread frontoparietal activation elicited by the 2-back task relative to the 1-back task.

However, activity in the 3-back task did not follow the hypothesized pattern. As mentioned previously, this non-linear load effect has been demonstrated in other fNIRS studies (Aghajani et al., 2017; Mandrick et al., 2013, 2016), and seems likely due to poor performance on the task. These results are consistent with the idea that when task demands exceed the current mental capacity of participants, they may disengage from the task and potentially, fail to recruit the necessary cognitive resources (Mandrick et al., 2013).

Interestingly, results of the PLS analysis that incorporated individuals’ accuracy by N-back level demonstrated evidence for an interaction of load and performance in the recruitment of the PFC. Specifically, this multivariate approach showed that changes in deoxyhemoglobin concentrations (HbR) in the medial SFG, which were only uncovered when examining the relationship between brain-behavior, but not when behavioral performance was not accounted for. Here, greater reductions in HbR (i.e., more activation) was positively related to performance on the 3-back task, unrelated to accuracy in the 2-back task, and negatively related to accuracy in the 1-back task. This pattern of results suggests that more automaticity during the 1-back task (i.e., less activation) led to better performance on this relatively easy task, and extensive recruitment of the PFC was required for high accuracy on the more difficult, cognitively demanding 3-back task.

This effect may reflect what has been proposed by the neural efficiency hypothesis: that participants with overall greater cognitive processing ability will show less activation during easy tasks and more during difficult tasks (Dunst et al., 2014; Neubauer & Fink, 2009). This is thought to result from the lower metabolic demands that a “more efficient” brain requires during cognitive tasks. Though the neural efficiency hypothesis is often framed as reflecting individual differences in intelligence, there is also evidence that this effect occurs as a result of more efficient strategies after adequate practice on a specific task (Sayala et al., 2006). Thus, one possibility for this interaction of task difficulty and prefrontal activation is that this reflects individual differences in the learning and adoption of effective strategies during practice. Interestingly, recent neuroimaging work has shown that individuals whose brains are in a more scale-free or fractal state tend to reap the benefits of practice to a greater degree than do those starting in a less scale-free state (Kardan, Layden, et al., 2020). Though scale-free neural dynamics have been demonstrated in fMRI and EEG (Churchill et al., 2016; Kardan, Layden, et al., 2020), whether this signal can be extracted from fNIRS data remains an open question.

The current study has a few limitations. Specifically, though it would have been ideal to get clean data from bilateral frontal and parietal cortex, the montage was only able to cover the right parietal cortex and the overall the signal-to-noise ratio (SNR) in these channels was very poor compared to the frontal optodes. By design, the Brain AnalyzIR Toolbox performs robust regression to downweight outliers and low SNR channels, which is necessary to avoid high false positive rates (Barker et al., 2013; Santosa et al., 2018). However, it is unclear whether the weaker effects in this area are due to lack of good signal (a false negative) or actually due to a lack of parietal cortical involvement in the N-back task relative to prefrontal cortex.

Additionally, it is unclear from these data why the PLS analysis found the load-dependent relationship in deoxyhemoglobin (HbR) concentration changes but not in oxyhemoglobin (HbO). In general, the HbO signal is larger than HbR which makes it easier to detect significant effects in task-based fNIRS, as is the case in this study’s N-back contrasts. Relative to HbO, the HbR signal is slower and more tightly coupled with the BOLD response of fMRI (Huppert et al., 2006). Therefore, one possibility is that the relationship between task performance and activation got stronger with increasing time performing the task, which may be reflected to a larger extent in the slower HbR signal. Additionally, as HbR is more sensitive to oxygen metabolism (and HbO is more sensitive to blood flow changes), it may be that this effect is more driven by metabolic changes. However, future investigations would be required to directly test these possibilities.

In conclusion, the present study demonstrates that fNIRS activation is sensitive to cognitive load and is differentially affected by performance based on task difficulty. This work demonstrates the efficacy of using robust statistical procedures to deal with unique statistical properties of fNIRS signals and the utility of implementing data-driven, multivariate techniques to elucidate more nuanced relationships between brain activity and behavior.

## Data & Code Availability Statement

fNIRS data, accuracy data, analysis code (for Brain AnalyzIR Toolbox, Behavioral PLS, and task accuracy), results, and n-back PsychoPy task code are all publicly available at: https://osf.io/sh2bf/

## Author Contribution Statement

K.L.M. and K.W.C. formulated the research question, designed the study, and developed the acquisition protocol. K.W.C. implemented the N-Back study. K.L.M. collected the data. K.L.M., C.C.I., and T.J.H. analyzed the data. M.G.B. supervised the project. K.L.M. wrote the first draft. K.W.C., C.C.I., T.J.H., & M.G.B. provided critical revisions. All authors approved the final version of the manuscript for submission.

## Acknowledgements

Supported by the National Science Foundation BCS-1632445 to M.G.B. the TFK Foundation and the John Templeton Foundation to M.G.B. (The University of Chicago Center for Practical Wisdom and the Virtue, Happiness and Meaning of Life Scholars group) the Grossman Institute for Neuroscience, Quantitative Biology and Human Behavior Shared Equipment Award to M.G.B.]; the Mansueto Institute for Urban Innovation [Postdoctoral Fellowship to K.W.C.].

Our fNIRS device was provided by the University of Chicago Grossman Institute for Neuroscience, Quantitative Biology, and Human Behavior (Shared Equipment Award).

We thank Jaime Young, Tanvi Laktahkia, Olivia Paraschos, and Anabella Pinton for their assistance in data collection.

## Footnotes

1 Three of the 62 usable fNIRS participants (#s P42, P67, and P70) were removed due to undue group-level leverage, see section *2.6.3. fNIRS Data Analysis: Pre-processing Pipeline and Task-Based Activation*.

